# Interaction between buprenorphine and norbuprenorphine in neonatal opioid withdrawal syndrome

**DOI:** 10.1101/2022.12.27.521115

**Authors:** Julia Tobacyk, Brian J Parks, Paloma Salazar, Lori U Coward, Michael D Berquist, Gregory S Gorman, Lisa K Brents

## Abstract

Buprenorphine (BUP) is the preferred treatment for opioid use disorder during pregnancy but can cause neonatal opioid withdrawal syndrome (NOWS). Norbuprenorphine (NorBUP), an active metabolite of BUP, is implicated in BUP-associated NOWS. We hypothesized that BUP, a low-efficacy agonist of mu opioid receptors, will not antagonize NorBUP, a high-efficacy agonist of mu opioid receptors, in producing NOWS. To test this hypothesis, we treated pregnant Long-Evans rats with BUP (0, 0.01, 0.1 or 1 mg/kg/day) ± NorBUP (1 mg/kg/day) from gestation day 9 until pup delivery, and tested pups for opioid dependence using our established NOWS model. We used LC-MS-MS to quantify brain concentrations of BUP, NorBUP, and their glucuronide conjugates. BUP had little effect on NorBUP-induced NOWS, with the exception of 1 mg/kg/day BUP significantly increasing NorBUP-induced NOWS by 58% in females. BUP and NorBUP brain concentrations predicted NOWS in multiple linear regression models. Interestingly, NorBUP contributed more to NOWS in females (β_NorBUP_ = 51.34, p = 0.0001) than in males (β_NorBUP_ = 19.21, P = 0.093), while BUP was similar for females (β_BUP_ = 10.62, P = 0.0017) and males (β_BUP_ = 11.38, P = 0.009). We are the first to report that NorBUP induces NOWS in the presence of BUP and that there is sexual divergence in the contribution of NorBUP to BUP-associated NOWS. These findings suggest that females are more susceptible to NorBUP-induced NOWS, and that treatment strategies that reduce prenatal NorBUP exposure may be more effective for females than males.

- Buprenorphine does not antagonize fetal norbuprenorphine dependence.
- In females, buprenorphine and norbuprenorphine have an additive effect.
- Norbuprenorphine contributes to NOWS in females more than in males.
- The glucuronide of norbuprenorphine accumulates in fetal brain.
- The glucuronide of buprenorphine does not accumulate in fetal brain.

## 1. Introduction

The United States (U.S.) is in the midst of an opioid addiction epidemic. According to the CDC, there were over 80,000 drug overdose deaths that involved opioids in the U.S. in 2021, which is the highest number ever recorded (CDC, 2022). The opioid crisis affects pregnant women and infants across the U.S., as evidenced by increasing rates of opioid use disorder (OUD) among pregnant women and rising rates of neonatal opioid withdrawal syndrome (NOWS) among infants (Corr and Hollenbeak, 2017; Patrick et al., 2015). Onset of NOWS occurs within the first days and weeks of life in newborns chronically exposed *in utero* to opioids, which induces physical opioid dependence in the fetus. The abrupt cessation of *in utero* opioid exposure at birth can lead to severe withdrawal symptoms, such as high-pitched crying, tremors, convulsions, vomiting, diarrhea, sleep disturbances and hypersensitivity to stimuli and often requires treatment in the neonatal intensive care unit with an average hospital stay of 16 days (Ramphul et al., 2020; Strahan et al., 2020). Diagnoses of maternal OUD at delivery and neonatal abstinence syndrome (NAS; a diagnosis that includes NOWS) each increased by more than 600% between 1999 and 2017 in the U.S. (Haight et al., 2018; Hirai et al., 2021; Patrick et al., 2012). Today, cases of NAS, which are driven primarily by prenatal opioid exposure, exceed even the most common birth defects, including cleft lip, cleft palate, clubfoot, and Down Syndrome combined (CDC). The clinical severity and high incidence of NOWS have driven the economic burden to treat NAS to approximately $2.5 billion per year (Ramphul et al., 2020).

Buprenorphine (BUP), a high-affinity, low-efficacy agonist of mu opioid receptors (Elkader and Sproule, 2005), is currently the gold standard treatment for OUD during pregnancy. BUP decreases illicit opioid use (Brogly et al., 2014) and decreases neonatal hospital stays (Zedler et al., 2016) when administered to pregnant women as part of a medication-assisted treatment program for OUD. However, ∼50% of newborns develop moderate-to-severe NOWS following prenatal BUP treatment (Jones et al., 2010). The assumed cause of BUP-associated NOWS is fetal exposure to the parent compound, BUP. However, neither maternal BUP dose nor umbilical cord plasma concentration at delivery correlate with NOWS severity (Shah et al., 2016; Wong et al., 2018), suggesting that metabolites of BUP mediate NOWS. Indeed, the concentration of the active metabolite NorBUP in human umbilical cord plasma positively correlates with NOWS severity (Shah et al., 2016). Compelling evidence from our lab and others suggests that fetal exposure to NorBUP, a high-affinity and high-efficacy agonist of mu opioid receptors (Huang et al., 2001), contributes to BUP-associated NOWS (Griffin et al., 2019; McPhie and Barr, 2000). However, interactions between BUP and NorBUP in causing NOWS is unknown. For example, BUP may plausibly antagonize NorBUP and thereby mitigate NorBUP-induced NOWS. Alternatively, NorBUP and BUP may have additive effects on NOWS, causing more severe NOWS to the neonate who was exposed to both during gestation. Studies that investigate the combined effects of BUP and NorBUP on the fetus are needed to predict the potential effectiveness of treatment strategies that reduce fetal exposure to NorBUP. Such studies will also define the importance of NorBUP in BUP-associated NOWS and will thus determine whether factors that enhance BUP metabolism to NorBUP (e.g., polymorphisms) increase susceptibility to NOWS.

The goal of this study was to determine interactions between BUP and NorBUP in causing NOWS. We hypothesized that BUP will not antagonize NorBUP-induced NOWS and that NorBUP co-administered with BUP will increase NOWS severity compared to administration of BUP alone. To test these hypotheses, we treated pregnant rats from gestation day (GD) 9 until delivery with BUP (0, 0.01, 0.1, or 1 mg/kg/day) in the presence or absence of a single dose of NorBUP (1 mg/kg/day) and tested their neonatal pups for withdrawal using our established model of NOWS. This approach enabled examination of BUP and NorBUP interactions by modeling a wide range of BUP: NorBUP dosing ratios (1:1, 10:1 and 100:1) and fetal brain exposures. The NorBUP dose (1 mg/kg/day) was selected because it produces blood concentrations in pregnant rats similar to those reported in pregnant women taking BUP for OUD (Concheiro et al., 2011; Griffin et al., 2019), thus improving the clinical relevance of the rat model. Because of previously reported sex differences in the risk of NOWS (Charles et al., 2017), we assessed withdrawal severity separately in male and female pups. To our knowledge, we are also the first to quantify concentrations of BUP, NorBUP, BUP glucuronide (BUP-gluc), and NorBUP glucuronide (NorBUP-gluc) in neonatal rat brains and relate these concentrations to NOWS severity. This study fills the knowledge gaps of the respective roles of BUP and NorBUP in BUP-associated NOWS. Examining the pharmacological profile of BUP and its metabolites in maternal-fetal dyads can enable the development of improved OUD treatments that exhibit lower NOWS incidence and severity.

## 2. Materials and Methods

### 2.1 Drugs and reagents

Buprenorphine HCl (BUP) and norbuprenorphine in free-base form (NorBUP) were provided by the National Institute on Drug Abuse (NIDA) Drug Supply Program (Bethesda, MD). Dimethyl sulfoxide (DMSO) and polyethylene glycol-400 (PEG-400) were purchased from Fisher Scientific. BUP, NorBUP, and the combination of BUP and NorBUP were dissolved in vehicle consisting of 1:2:1 DMSO/PEG-400/sterile saline and loaded into Alzet 2ML2 osmotic minipumps (Cupertino, CA). Prior to surgical subcutaneous implantation, osmotic minipumps were incubated in sterile saline at 37°C overnight as per manufacturer’s instructions. Alzet model 2ML2 osmotic minipumps deliver their contents at a constant rate of 0.120 mL per day for 14 days. BUP was formulated to deliver 0.01, 0.1, or 1 mg/kg/day and NorBUP was formulated to deliver 1 mg/kg/day. The concentrations of drug formulations were based on GD 8 weights (a day prior to osmotic minipump implantation). Naltrexone hydrochloride (NTX) was purchased from Sigma-Aldrich (St. Louis, MO) and dissolved in sterile saline prior to administering via the intraperitoneal (i.p.) route to rat pups.

With the exception of terfenadine, all drugs and standards used for LC/MS/MS analyses of neonatal brain tissues were purchased from Sigma-Aldrich (St. Louis, MO). Terfenadine was purchased from Fisher Scientific (Atlanta, GA). Buprenorphine HCl and norbuprenorphine were purchased as 1 mg/mL solutions in methanol. Buprenorphine-3-beta-D-glucuronide, norbuprenorphine glucuronide, and the internal standards buprenorphine-D_4_-3-beta-D-glucuronide and norbuprenorphine glucuronide-D^3^ were purchased as 100 μg/mL solutions in methanol.

For the [^35^S]GTPγS binding assay, [^35^S]GTPγS (1250 Ci/mM) was purchased from PerkinElmer (Waltham, MA). Guanosine-5-diphosphate (GDP) and guanosine 5’-O-[gamma-thio]triphosphate (GTPγS) were purchased from Fisher Scientific and Sigma-Aldrich, respectively.

### 2.2 Animal care and use

This study was performed in accordance with the Guide for the Care and Use of Laboratory Animals. All animal protocols were approved by Institutional Animal Care and Use Committee at University of Arkansas for Medical Sciences (UAMS) prior to commencement of experiments.

Fifty-six timed-pregnant Long-Evans rats (Charles River Laboratories, Wilmington, MA) arrived at the UAMS animal facility on GD 5 or 6 weighing between 175–230 g. Pregnant rats were singly housed in Plexiglas cages (30 cm x 24.6 cm x 41.2 cm) with environmental enrichment (crinkled paper and a plastic tunnel) and had *ad libitum* access to food and water. The Allentown NexGen Rat 900 Ecoflo system was used to ventilate each cage.

On GD 9, osmotic minipumps containing BUP, NorBUP, or the combination of BUP and NorBUP or vehicle control (1:2:1 DMSO:PEG400:saline) were subcutaneously implanted per manufacturer’s guidelines to deliver one of the following eight treatments: 1) vehicle control, 1:2:1 DMSO:PEG-400:sterile saline; 2) 1 mg/kg/day NorBUP; 3) 0.01 mg/kg/day BUP; 4) 0.1 mg/kg/day BUP; 5) 1 mg/kg/day BUP; 6) 0.01 mg/kg/day BUP + 1 mg/kg/day NorBUP; 7) 0.1 mg/kg/day BUP + 1 mg/kg/day NorBUP; and 8) 1 mg/kg/day BUP + 1 mg/kg/day NorBUP (**Fig. 1**). The study was performed using four cohorts of rats in the following order: cohort 1 (n=8), cohort 2 (n=16), cohort 3 (n=16) and cohort 4 (n=16). Treatment assignments were balanced such that each cohort had either n=1 (cohort 1) or n=2 (cohorts 2-4) dams for each treatment, yielding n=7 per treatment (**Table 1**). Pregnant dams were left undisturbed, except for daily health checks, weekly cage changes, and weight measurements taken on weekdays. Neonatal pups were separated from the dam three to twelve hours after delivery and tested immediately as described in “Neonatal opioid withdrawal testing” below. Dams were euthanized in accordance with guidelines from the American Veterinary Medical Association within 24 hr of precipitated withdrawal pup testing. Maternal plasma and remaining minipump content were collected and stored in at -80°C. Spontaneous abortions were assessed as lack of presence of pups or signs of recent delivery (e.g., blood, placenta) by end of gestation day 23 *and* early weight gain rates that were equal to those of dams that successfully delivered pups, but plateaued following minipump implantation, suggesting the dam was pregnant but spontaneously aborted.

**Table 1.**

|  | Vehicle | BUP<br>0.01 | BUP<br>0.1 | BUP 1 | BUP<br>0.01 +<br>NorBUP<br>1 | BUP 0.1<br>+<br>NorBUP<br>1 | BUP 1 +<br>NorBUP<br>1 | NorBUP<br>1 |
| --- | --- | --- | --- | --- | --- | --- | --- | --- |
| *N* (number of<br>dams that<br>delivered out<br>of 7) | 6 | 6 | 6 | 4 | 6 | 7 | 7 | 7 |
| Spontaneous<br>Abortions, % | 14.3 | 14.3 | 14.3 | 42.9 | 14.3 | 0 | 0 | 0 |
| Litter size,<br>mean (S.D.) | 12.5<br>(3.3) | 12.0<br>(1.3) | 11.0<br>(2.0) | 11.8<br>(1.3) | 12.3<br>(2.3) | 10.6<br>(2.7) | 12.9<br>(2.0) | 11.7<br>(3.7) |
| Males per<br>litter, mean<br>(S.D.) | 6.5<br>(1.6) | 5.3<br>(1.6) | 5.8<br>(2.1) | 6.0<br>(1.4) | 5.8 (1.7) | 4.1 (1.6) | 5.9 (0.7) | 7.4 (1.8) |
| Females per<br>litter, mean<br>(S.D.) | 6.0<br>(2.4) | 6.7<br>(0.8) | 5.2<br>(2.2) | 5.8<br>(0.5) | 6.5 (1.9) | 7.1 (2.7) | 8.7 (3.1) | 7.7 (3.7) |
| Sex ratio,<br>males :<br>females | 1.2<br>(0.4) | 0.8<br>(0.3) | 1.4<br>(1.0) | 1.1<br>(0.3) | 1.0 (0.7) | 0.7 (0.3) | 0.7 (0.3) | 1.2 (1.0) |
| Neonatal<br>male pup<br>weights, g,<br>mean (S.D.) | 6.2<br>(0.4) | 5.7<br>(1.2) | 6.6<br>(0.5) | 6.5<br>(0.2) | 6.4 (0.7) | 7.1 (1.0) | 6.3 (0.7) | 6.1 (0.9) |
| Neonatal<br>female pup<br>weights, g,<br>mean (S.D.) | 6.0<br>(0.4) | 6.1<br>(0.4) | 6.4<br>(0.5) | 6.0<br>(0.5) | 6.0 (0.6) | 6.0 (1.7) | 6.0 (0.7) | 6.0 (0.5) |

**Figure 1.**
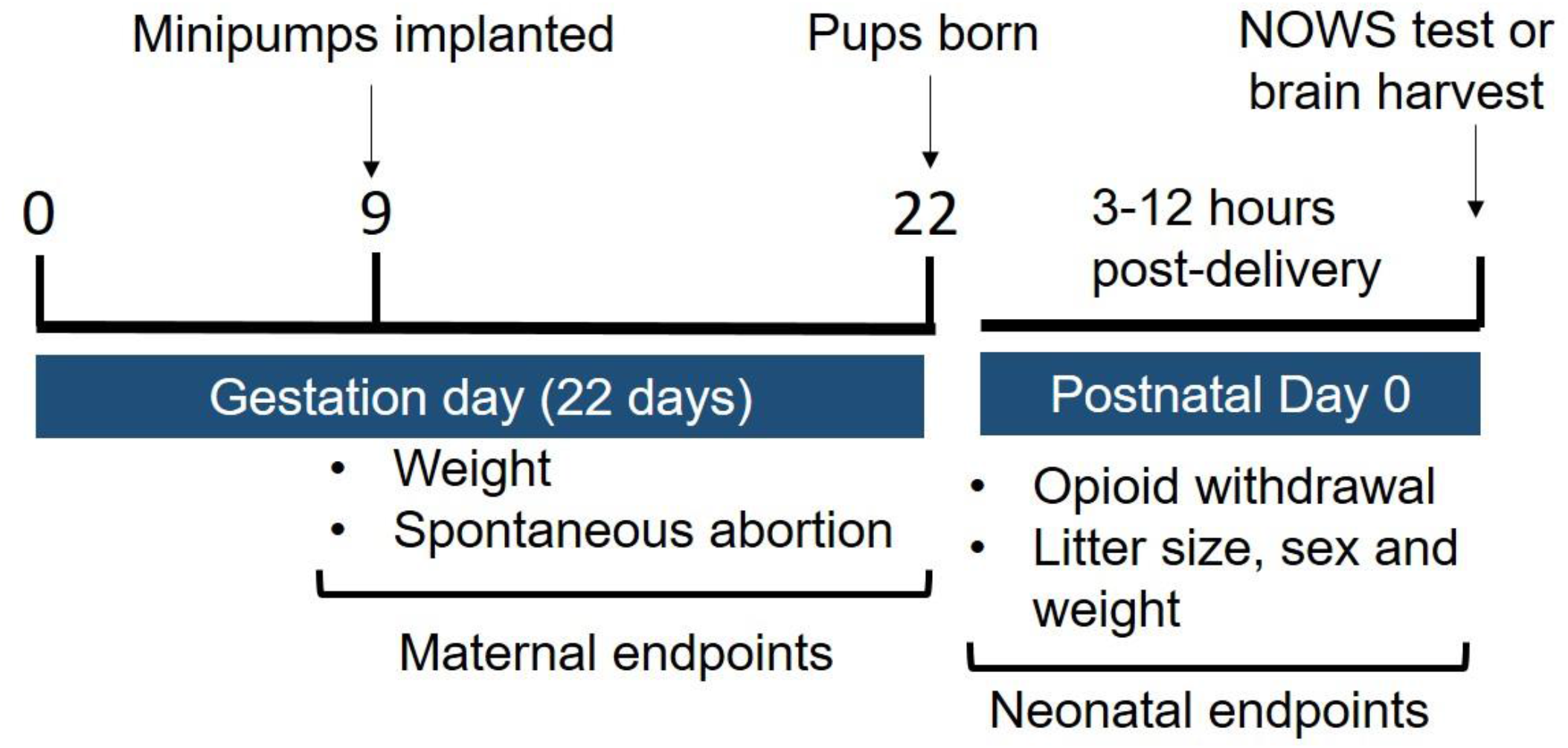
Experimental design. GD 5-6 pregnant rats arrived to the animal facility. On GD 9, osmotic minipumps with BUP (0, 0.01, 0.1 and 1 mg/kg/day) and/or NorBUP (0 and 1 mg/kg/day) were subcutaneously implanted in pregnant rats. Maternal weight gain and spontaneous abortion were monitored throughout gestation. Neonatal opioid withdrawal and litter characteristics were measured immediately following separation from dam, which was 3-12 hours after delivery. Neonatal brains of pups that did not undergo withdrawal testing were collected during withdrawal testing to quantify concentrations of BUP, NorBUP, and their glucuronide conjugates.

### 2.3 Neonatal opioid withdrawal testing

Two hundred and ninety-four neonatal rat pups from a total of 49 litters were tested for NTX-precipitated opioid withdrawal. Upon separation from their dams, rat pups were counted, grouped by sex, and weighed. A total of six pups (three females and three males) from each litter were randomly selected and administered (i.p.) one of the following treatments: saline (10 mL/kg), 1 mg/kg NTX, or 10 mg/kg NTX to precipitate withdrawals. Saline was administered as a postnatal challenge to serve as a control for, and assess the effects of, injection and spontaneous withdrawal on pup behaviors. As we previously described (Griffin et al., 2019), pups were placed in test chambers (12.86 cm x 10.16 cm x 3.00 cm) containing aspen bedding and videoed for 10 minutes beginning immediately after injection. Investigators conducting the testing were blinded to prenatal pup treatments. Pup movements (as “movement duration” in seconds) were quantified in real time using Noldus Ethovision XT automated software (v. 15) using previously validated methods (Kulbeth et al., 2021), with minor modifications of increasing sample rate from 5 samples/second to 30 samples/second and applying the default built-in image processing algorithm for data smoothing to remove noise (Kulbeth et al., 2021). Threshold for start and stop velocities for “movement” were set at 0.4 and 0.3 cm/s, respectively. “Gray scaling” was used as the detection method and detection setting were optimized for each cohort separately. At the final step of data acquisition, playback control and the Video Time Slider within Ethovision XT was used in a quality control step to confirm that pup movements were detected with minimal noise for the entire 10 min testing session. Pups not undergoing withdrawal testing were sacrificed within 10 min of maternal separation for brain tissue harvest. Brains were snap-frozen in liquid nitrogen, stored at -80°C and analyzed by LC/MS/MS for the following metabolites: BUP, NorBUP, BUP glucuronide, and NorBUP glucuronide concentrations (see “Quantitative measurement of BUP, NorBUP, and glucuronide conjugates of BUP and NorBUP in neonatal brains” below). Pups tested for withdrawals were euthanized immediately after withdrawal testing. All pups were euthanized using methods in accordance with the American Veterinary Medical Association’s guidelines.

### 2.4 Quantitative measurement of BUP, NorBUP, and glucuronide conjugates of BUP and NorBUP in neonatal brains

Neonatal rat brain samples were homogenized 1:5 with 5 mM ammonium acetate buffer and spiked with internal standards (20 ng/mL terfenadine and 50 ng/mL buprenorphine-D_4_-3-beta-D-glucuronide and norbuprenorphine glucuronide-D_3_ in acetonitrile). Blanks, calibration standards, and quality control samples were made by spiking brain homogenates from untreated control neonatal rats with known amounts of analytes and internal standards. For each analyte, calibration standards ranged from 0.05 to 50 ng/mL of homogenate (0.25 to 250 ng/g of brain), and quality control samples contained 0.2, 2, or 20 ng/mL (1, 10, or 100 ng/g) of each analyte. Acetonitrile (1 mL) was added to each sample to precipitate the protein. Samples were vortexed and centrifuged at 21,000x*g* for five minutes. Supernatant was transferred to a clean test tube, and acetonitrile was evaporated from the supernatant under nitrogen in a 50°C water bath. The remaining residue was suspended in 50/50 5 mM ammonium acetate/acetonitrile (200 μL) and transferred to a limited volume autosampler vial and analyzed by HPLC/Mass Spectrometry/Mass Spectrometry (HPLC/MS/MS). The LC/MS/MS system consisted of Shimadzu system (Columbia MD) equipped with LC20-AD dual HPLC pumps, an SIL20-AC HT autosampler, and a DGU-20A2 inline degasser. Detection was performed using an Applied BioSystems 5500 QTRAP (Applied Biosystems, Foster City, CA) triple quadrupole mass spectrometer operated in the positive ion mode. Mass calibration, data acquisition, and quantitation were performed using Applied Biosystem Analyst 1.7.2 software (Applied Biosystems, Foster City, CA). For each analyte, the lower limit of quantitation (LLOQ) was 0.05 ng/mL (0.25 ng/g) and the upper limit of quantitation (ULOQ) was 50 ng/mL (250 ng/g).

### 2.5 Cell Culture

Chinese hamster ovary (CHO) cells transfected with human mu opioid receptors (CHO-hMOR) were provided by Dr. Dana Selley (Virginia Commonwealth University, Richmond, VA). CHO-hMOR cells were cultured in DMEM Nutrient Mix F12 1:1 media containing 10% FetalPlex®, 1% penicillin-streptomycin, and 0.2 mg/mL hygromycin (Gibco, Waltham, MA). CHO-hMOR were grown in sterile T175 flasks and kept in a humidified chamber with 5% CO_2_ at 37°C. Cells were harvested at 100% confluence with 0.04% EDTA in phosphate buffered saline (pH 7.5) and centrifuged at 1040 *x g* to obtain a cell pellet. Supernatant was discarded and cell pellets were stored at -80°C until they were used to make membrane homogenates for the [^35^S]GTPγS assay.

### 2.6 Membrane homogenates preparation

CHO-hMOR pellets were suspended in 10 mL (or ∼2 mL/cell pellet) of cold homogenization buffer (50 mM HEPES, pH 7.4, 3 mM MgCl_2_, and 1 mM EGTA). A 40 mL Dounce glass homogenizer was used to homogenize the cell membranes. The homogenate was centrifuged at 40,000 *x g* for 10 minutes at 4°C. The supernatant was discarded and the pellet was again transferred to the homogenizer, resuspended in 10 mL of homogenization buffer, coarsely homogenized and centrifuged at 40,000 *x g* for 10 minutes at 4°C. This step was repeated twice. The final pellet was homogenized using 10 strokes of a fine-grade pestle in 10 mL (or ∼2 mL/cell pellet) of cold 50 mM HEPES. Protein concentrations of homogenates were determined using the BCA Protein Assay (Thermo Scientific, Walham, MA). Homogenates were stored at -80°C until they were used in [^35^S]GTPγS binding assays.

### 2.7 [^35^S]GTPyS binding assay

Interactions between BUP and NorBUP-induced G-protein activation via hMOR were assessed by incubating the following in a 1-mL volume of buffer composed of 20 mM HEPES, 10 mM MgCl_2_, 100 mM NaCl, and 0.05% bovine serum albumin at 30°C for 30 minutes: 0.1 nM [^35^S]GTPγS; 50 μg CHO-hMOR membrane homogenates; varying concentrations of BUP ranging from 30 to 1000 nM or vehicle (0.01% DMSO); NorBUP (1,000 nM) or vehicle (0.01% DMSO); and 10,000 nM GDP. A subset of samples intended to quantify non-specific binding of the radioligand contained a high concentration of unlabeled GTPγS (10,000 nM) instead of BUP or NorBUP. Homogenates were pre-incubated with GDP for 5 mins prior to addition of all other components. Samples were conducted in triplicate in three independent experiments. Reactions were terminated by addition of 1 mL ice cold buffer (500 mM HEPES, 0.05% bovine serum albumin) and rapid filtration by a Brandel 24-sample cell harvester onto Whatman GF/B glass fiber filters. Sample tubes and filters were rinsed with five volumes of the ice-cold buffer and immediately placed in 7 mL scintillation vials containing 4 mL of ScintiVerse™ BD Cocktail scintillation fluid. Following an overnight incubation in the scintillation fluid, decays per minute (DPMs) of radioactivity in each sample was read by a Tri Carb 4910 TR Liquid Scintillation Counter (PerkinElmer, Waltham, MA). Specific binding of [^35^S]GTPγS to G-proteins was calculated for each sample by subtracting the number of DPMs quantified in non-specific binding samples (i.e. those containing only unlabeled GTPγS) from the number of DPMs quantified in each sample.

### 2.8 Statistical Analysis

All data are reported as mean ± S.E.M, unless otherwise stated. All analyses are performed using GraphPad Prism, version 9.4.1. (La Jolla, CA).

The main and interaction effects of BUP dose (0, 0.01, 0.1, or 1 mg/kg/day) and NorBUP dose (0 or 1 mg/kg/day) on neonatal withdrawal (as measured by movement duration in seconds) were evaluated using a two-way ANOVA analysis within each sex and postnatal challenge group (0, 1, and 10 mg/kg NTX). Šidák’s multiple comparisons test was used to identify the simple effect of NorBUP at each BUP dose and thus determine whether the presence of NorBUP affects BUP-induced withdrawal. Dunnett’s multiple comparisons test was used to compare each BUP dose in the presence of NorBUP to NorBUP alone and thus determine whether the presence of BUP affects NorBUP-induced withdrawal.

To determine whether neonatal withdrawal differed between males and females, we used a two-way ANOVA, with neonatal withdrawal (as measured by movement duration in seconds) as the dependent variable and prenatal treatment group (8 levels) and sex (2 levels) as the independent variables. Šidák’s multiple comparison tests were conducted to identify simple effects of sex within each treatment group. Separate analyses were conducted for each postnatal challenge group (0, 1 and 10 mg/kg NTX).

To identify potential confounders and better understand the prenatal effects of the drug treatments, we assessed multiple variables of maternal and neonatal health. Maternal gestational weight gain was measured at least five times a week between GD 7 and GD 21. Group-wise linear regression analyses were performed on weight data from each dam that delivered pups (i.e., weights of dams that did not deliver were excluded). An F-test was used to test for statistical differences in slope estimates between regression lines. The relationship between presence of NorBUP and probability of delivering was assessed using Fisher’s exact test. Litter characteristics (mean litter size, mean female and male pup weights, male: female ratio, and mean number of female and male pups per litter) were assessed using one-way ANOVA at the treatment group level (8 levels) followed by Dunnett’s multiple comparisons test.

The main and interaction effects of BUP dose (0, 0.01, 0.1, or 1 mg/kg/day) and NorBUP dose (0 or 1 mg/kg/day) on BUP, NorBUP, BUP-gluc and NorBUP-gluc metabolites (as measured in neonatal male and female brains in ng/g) were evaluated using a two-way ANOVA analysis in the 1 mg/kg NTX postnatal challenge group. Šidák’s multiple comparison test was used to identify the simple effect of NorBUP at each BUP dose.

Multiple linear regression analysis was used to determine the relationship between BUP metabolite concentration (BUP, NorBUP, BUP-gluc and NorBUP-gluc) and neonatal withdrawal (as measured by movement duration in seconds) both in males and females. Neonatal brain concentrations and movement duration measurements were matched from same-sex littermates, with the assumption that the pooled brain concentrations that were measured from littermates not undergoing NOWS testing closely represented those of the pups undergoing NOWS testing whose brains were not measured.

For the [^35^S]GTPyS binding assay, the main and interaction effects of BUP concentrations and NorBUP concentration on specific binding measured in DPMs were evaluated using a two-way ANOVA analysis with repeated measures (sphericity assumed). Šidák’s multiple comparison test was used to identify the simple effect of NorBUP (0 or 1000 nM) at each BUP (0, 3, 10, 30, 100, 300, and 1000 nM) concentration and Dunnett’s multiple comparison test was used to compare each BUP concentration in the presence of NorBUP to NorBUP alone and thus determine whether the presence of BUP affects NorBUP-induced withdrawal. Statistical significance was declared at *p* <0.05 for all analyses.

## 3. Results

Neonatal withdrawal was quantified as movement duration in seconds following prenatal treatment with BUP (0, 0.01, 0.1 or 1 mg/kg/day) in the absence or presence of NorBUP (0 or 1 mg/kg/day) following postnatal challenge with saline, 1 mg/kg NTX or 10 mg/kg NTX to precipitate withdrawal. There was no interaction between BUP and NorBUP in males or females following any postnatal challenge (**Fig 2**). Following postnatal challenge with saline, there was a main effect of BUP for males and females (**males**: F [3, 41] = 14.85, \*\*\*\**p<*0.0001; and **females**: F [3, 41] = 23.88 \*\*\*\**p<*0.0001), and no main effect of NorBUP (**males**: F [1, 41] = 0.01290, *p=*0.9101 and **females**: F [1, 41] = 0.01387; *p=*0.9068; **Fig 2A and 2D**). In males and females, the highest dose of BUP (1 mg/kg/day) increased neonatal movement duration relative to vehicle control following postnatal challenge with saline (**Fig 2A and 2D; males**: ^##^*p=*0.0016 and **females**: ^###^*p<*0.0001, Dunnett’s multiple comparisons test). In the presence of NorBUP (1 mg/kg/day), the highest dose of BUP (1 mg/kg/day) significantly increased neonatal movement duration relative to NorBUP alone (**Fig 2A and 2D; males:** ^###^*p=*0.0007 and **females**: ^###^*p=*0.0004, Dunnett’s multiple comparisons test). The presence of NorBUP had no significant effect at any dose of BUP (P>0.05, Šidák’s multiple comparison test comparing presence vs. absence of NorBUP for each BUP dose; **Fig 2A and 2D**).

**Figure 2.**
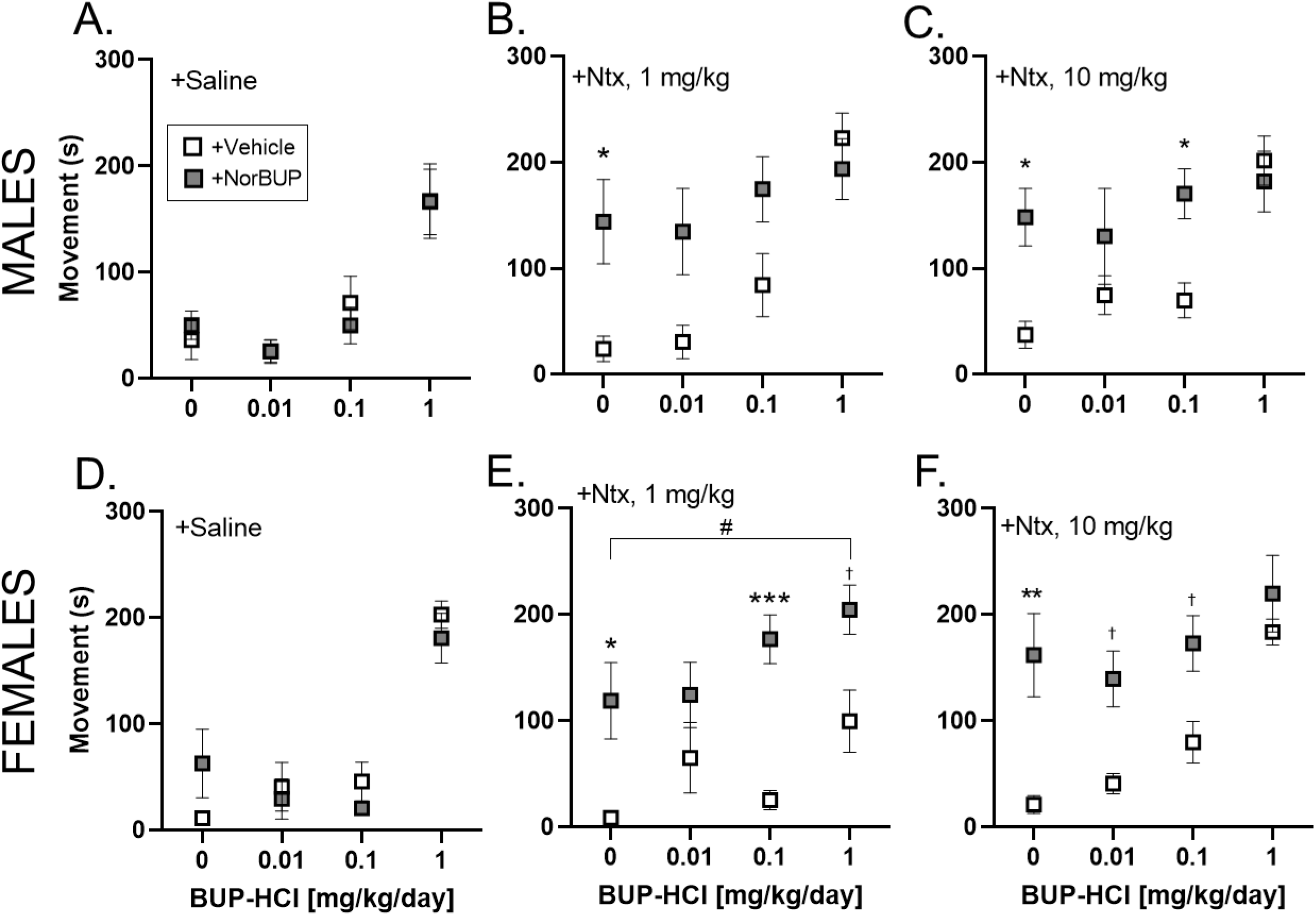
Neonatal withdrawal signs differ in males and females. Quantification of neonatal withdrawal signs in males (A-C) and females (D-F) following prenatal treatment with BUP (0, 0.01, 0.1 and 1 mg/kg/day), NorBUP (0 and 1 mg/kg/day) and the combination of both BUP and NorBUP. Neonatal pups were challenged with NTX (0, 1 and 10 mg/kg, i.p.). Points represent group means and error bars represent S.E.M; n=4-7. ^†^P < 0.10, **P < 0.01, ***P < 0.001, ****P < 0.0001 vs BUP + vehicle, Šidák’s multiple comparisons test. ^&^P < 0.10, ^#^P < 0.05, ^##^P < 0.01, ^###^P < 0.001 vs NorBUP + vehicle, Dunnett’s multiple comparison test.

Following postnatal challenge with the intermediate dose of NTX, 1 mg/kg, there were significant main effects of BUP and NorBUP for males and females (**males**: BUP main effect (F [3, 41] = 6.586, ***p=.0010); NorBUP main effect (F [1, 41] = 10.54, **p=.0023); **females**: BUP main effect (F [3, 41] = 3.525, *p=0.0231); NorBUP main effect (F [1, 41] = 31.78, \*\*\*\**p<*0.0001); **Fig 2B and 2E**).

For males challenged with 1 mg/kg NTX (**Fig 2B)**, NorBUP significantly increased withdrawal relative to vehicle control (\**p=*0.0273, Šidák’s multiple comparison test), but not in the presence of any BUP dose relative to the BUP dose alone (P = 0.0859 with 0.01 mg/kg/day BUP; P = 0.1423 with 0.1 mg/kg/day BUP; P = 0.9561 with 1 mg/kg/day BUP, Šidák’s multiple comparison test). Co-administration of NorBUP and BUP did not significantly increase or decrease withdrawal relative to NorBUP alone (P > 0.48 in the presence of each BUP dose, Dunnett’s multiple comparison test). The highest dose of BUP (1 mg/kg/day) significantly increased withdrawal relative to vehicle control (^###^*p=*0.0006, Dunnett’s multiple comparison test). The lower two doses of BUP (0.01 and 0.1 mg/kg/day) did not significantly increase withdrawal relative to vehicle control (P > 0.39, Dunnett’s multiple comparison test).

For females that were challenged with 1 mg/kg NTX, NorBUP increased withdrawal relative to vehicle control (\**p=*0.016) and in the presence of 0.1 mg/kg/day BUP (\*\*\**p=*0.0006) and 1 mg/kg/day BUP (^†^P =0.054) relative to 0.1 and 1 mg/kg/day of BUP alone, respectively (Šidák’s multiple comparison test; **Fig 2E**). Co-administration of NorBUP and the highest dose of BUP (1 mg/kg/day) significantly increased withdrawal by 58% relative to NorBUP alone (^#^*p=*0.048, Dunnett’s multiple comparison test; **Fig 2E**). No dose of BUP, in the absence of NorBUP, significantly increased withdrawal relative to vehicle control (P > 0.09, Dunnett’s multiple comparison test).

Following challenge with the highest dose of NTX (10 mg/kg) there were significant main effects of BUP and NorBUP for males and females (**males**: BUP main effect (F [3, 41] = 4.943, \*\**p=*0.0051); NorBUP main effect (F [1, 41] = 10.36; \*\**p=*0.0025); **females**: BUP main effect (F [3, 41] = 6.508, \*\**p=*0.0011); NorBUP main effect (F [1, 41] = 22.09, \*\*\*\**p<*0.0001); **Fig 2C and 2F**).

For males challenged with 10 mg/kg NTX (**Fig 2C**), NorBUP significantly increased withdrawal relative to vehicle control (\**p=*0.018) and in the presence of 0.1 mg/kg/day BUP relative to 0.1 mg/kg/day of BUP alone (\**p=*0.037, Šidák’s multiple comparison test; **Fig 2C**). Co-administration of NorBUP and BUP did not significantly increase or decrease withdrawal relative to NorBUP alone (P>0.66) in the presence of each BUP dose, Dunnett’s multiple comparison test). In the absence of NorBUP, the lower two doses of BUP (0.01 and 0.1 mg/kg/day) had no significant effect on withdrawal relative to vehicle control (P > 0.65, Dunnett’s multiple comparison test); however, the highest dose of BUP (1 mg/kg/day) significantly increased withdrawal (^##^*p=*0.0012, Dunnett’s multiple comparison test; **Fig 2C**).

For females that were challenged with 10 mg/kg NTX, NorBUP significantly increased withdrawal relative to vehicle control (\*\**p=*0.002) and in the presence of 0.01 mg/kg/day BUP (^†^*p=*0.061) and 0.1 mg/kg/day BUP (^†^*p=*0.069) relative to 0.01 and 0.1 mg/kg/day of BUP alone (Šidák’s multiple comparison test; **Fig 2F**). Co-administration of NorBUP and BUP did not significantly increase or decrease withdrawal relative to NorBUP alone (P > 0.26 in the presence of each BUP dose, Dunnett’s multiple comparison test). In the absence of NorBUP, the lower two doses of BUP (0.01 and 0.1 mg/kg/day) had no significant effect on withdrawal relative to vehicle control (P > 0.32, Dunnett’s multiple comparison test); however, the highest dose of BUP (1 mg/kg/day) significantly increased withdrawal (^##^*p=*0.0017, Dunnett’s multiple comparison test; **Fig 2F**).

We assessed for sex effects by testing for main and interaction effects of sex and treatment group membership on movement duration, and by testing for sex differences within each treatment group (**Fig 3**). These analyses were performed separately within each NTX-challenge group. There were no interactions between sex and treatment group membership and no main effects of sex for any NTX-challenge. As expected, there was a main effect of treatment group within each of the NTX challenges (**saline:** F [7, 82] = 17.25; *p<*0.0001; **1 mg/kg NTX:** F [7, 82] = 11.11; *p<*0.0001; and **10 mg/kg NTX**: F [7, 82] = 11.34; *p<*0.0001; **Fig 3A-C**). There was no difference between males and females in opioid withdrawal within any treatment group (P≥ 0.118; Šidák’s multiple comparison test; **Fig 3**).

**Figure 3.**
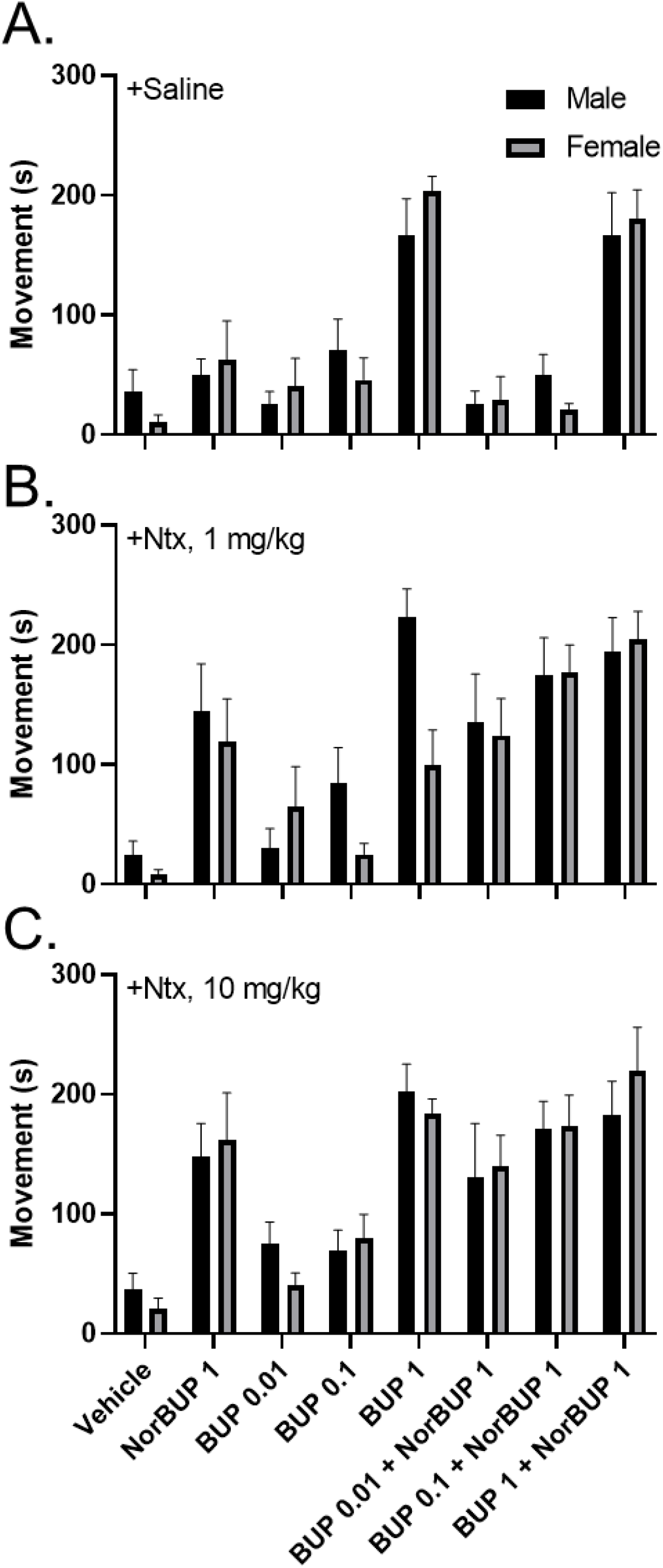
No sex differences were observed in NOWS. Sex effects of prenatal treatment on neonatal withdrawal. Bars represent the mean movement (y-axis) of pups grouped by sex (male, black bars; female, grey bars) that were postnatally challenged with saline (A), 1 mg/kg Ntx (B) and 10 mg/kg Ntx (C). Prenatal treatments are plotted on the x-axis and include NorBUP (1 mg/kg/day; s.c.), BUP (0.01, 0.1 and 1 mg/kg/day; s.c.) and vehicle (1:2:1 DMSO/PEG-400/saline, 0.120 mL/day, s.c.). Error bars represent S.E.M.

We assessed several maternal and neonatal health endpoints to identify potential confounding effects of the prenatal treatments on neonatal opioid withdrawal testing and maternal-fetal opioid pharmacokinetics, including poor maternal-fetal nutrition, poor weight gain, and sex-specific and non-sex-specific fetal absorptions. These endpoints were maternal weight gain (**Fig 4**), spontaneous abortion rate, litter size, sex distribution, and pup weights (**Table 1**). Slow rate of maternal weight gain suggests reduced food consumption and/or poor maternal/fetal health, which may affect pup size and movement. Maternal weight gain did not differ among the groups (F [7, 41] = 0.2935, *p=*0.9564, pooled slope = 8.288 g/day; **Fig 4**). Weights of the dams that did not deliver pups (n = 7) were excluded from this analysis. Early in gestation (GD 6-9), the dams that did not deliver pups gained weight at the same rate as dams that delivered (P = 0.1471), but their weight gain slowed after osmotic minipump implantation on GD 9 relative to the dams that eventually delivered pups (*p<*0.0001). This suggests that these dams experienced spontaneous abortions instead of unsuccessful fertilization and implantation.

**Figure 4.**
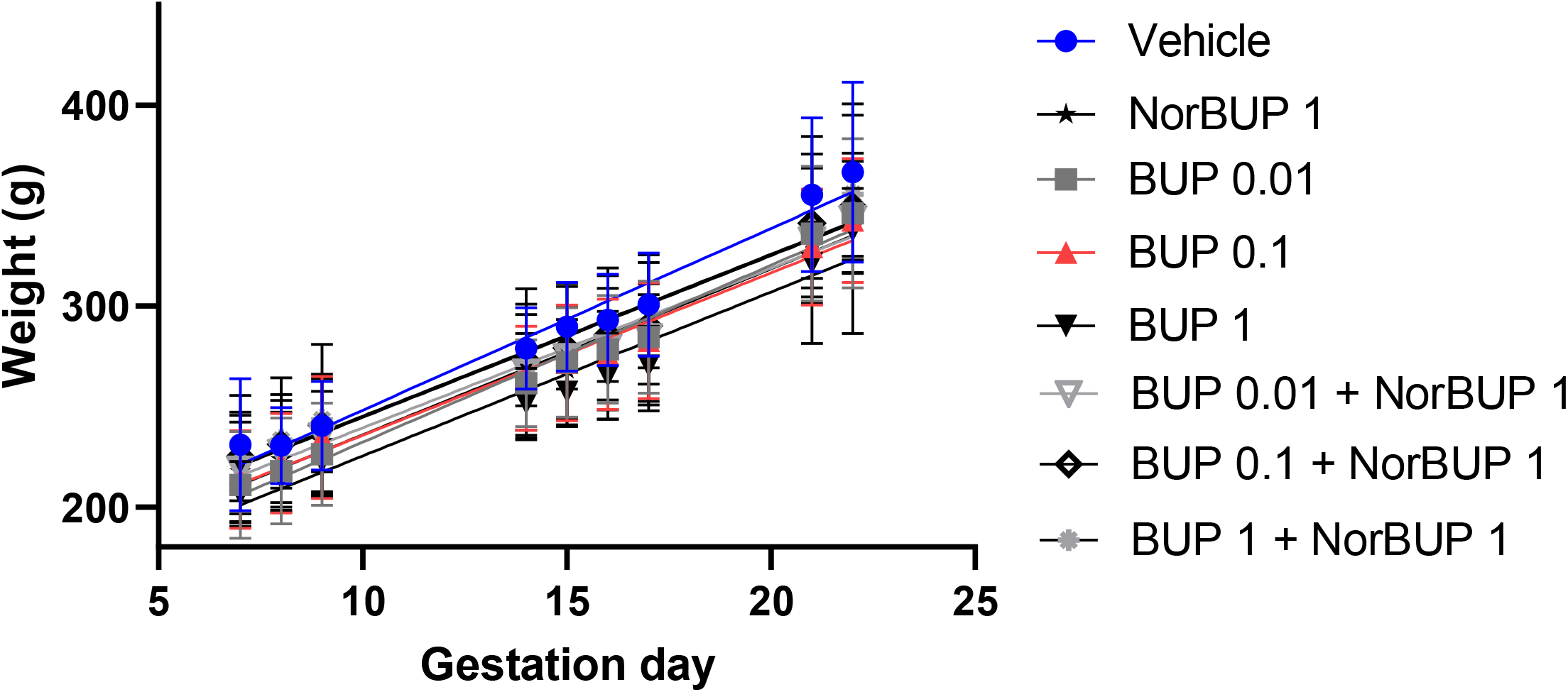
Maternal weight gain was not affected by prenatal treatment. Data points represent group mean weights (y-axis) plotted by gestation day (x-axis); error bars represent S.E.M. Each line represents a linear regression analysis best-fit of the data points for each prenatal treatment group. There were no group differences in the slopes of the linear regression (F[7,41] = 0.2935, *p=*0.9564), pooled slope = 8.288 g/day.

Spontaneous abortions were relatively rare (≤1 out of 7 dams/group) except in the group treated with 1 mg/kg/day BUP (42.9%). Because of the small group sizes, we were unable to perform analyses to determine whether drug treatment affected probability of delivery. However, we assessed whether NorBUP affected the probability of delivery by performing Fisher’s exact test on the data grouped by presence or absence of NorBUP (ignoring BUP dose). We determined that NorBUP had no significant effect on probability of delivery (*p=*0.101; relative risk = 0.81, 95% CI = 0.62 to 0.99). One-way ANOVAs showed no effect of treatment on litter size (F [7, 41] = 0.6024, *p=*0.7503), pup weight (**males**: F [7, 41] = 1.649, *p=*0.1492; **females**: F [7, 41] = 0.2166, *p=*0.9795) or sex ratio within (F [7, 41] = 1.165, *p=*0.3436). The number of females per litter was not affected by treatment (F [7, 41] = 1.278, *p=*0.2849). However, the number of males per litter was affected by prenatal treatment (F [7, 41] = 2.283, \**p=*0.0464). This effect was likely driven by the group that was treated with 0.1 mg/kg/day BUP + 1 mg/kg/day NorBUP, which averaged 2.4 fewer pups per litter than the vehicle control group (^†^*p=*0.065, Dunnett’s multiple comparison test).

Next, we quantified concentrations of BUP, NorBUP, and their glucuronide conjugates BUP-gluc and NorBUP-gluc in neonatal brain samples that had been pooled by litter and sex. These analyses enable us to assess how maternal dosing of BUP and NorBUP affects fetal/neonatal brain exposure to these opioids, and how fetal/neonatal brain exposures, in turn, affect neonatal opioid dependence and withdrawal. As expected, increasing maternal BUP dose increased neonatal brain concentrations of BUP, regardless of the presence or absence of NorBUP (**Fig 5A-B**). However, this increase was not linear. Increasing BUP dose by 10-fold (from 0.01 to 0.1 mg/kg/day and from 0.1 to 1 mg/kg/day) increased neonatal brain concentrations by only 3.0- to 4.6-fold, and increasing maternal BUP dose by 100-fold (from 0.01 to 1 mg/kg/day) increased neonatal brain concentrations by only 12.4- to 17-fold. Neonatal brain concentrations of BUP-Gluc were below the limits of quantitation (0.25 ng/g) for all but the highest dose of BUP (**Fig 5C-D**). For the highest dose of BUP, neonatal brain concentrations of BUP-Gluc were less than 1/13th those of BUP, suggesting maternal dosing with BUP causes very little fetal brain exposure to BUP-Gluc. Equal doses of BUP and NorBUP (1 mg/kg/day of each) yielded approximately 3-fold higher brain concentrations of BUP relative to NorBUP (compare **Fig 5A-B** to **Fig 5E-F**). Brains from pups whose dams were treated with NorBUP had approximately 3- to 4-fold higher concentrations of NorBUP-gluc than NorBUP.

**Figure 5.**
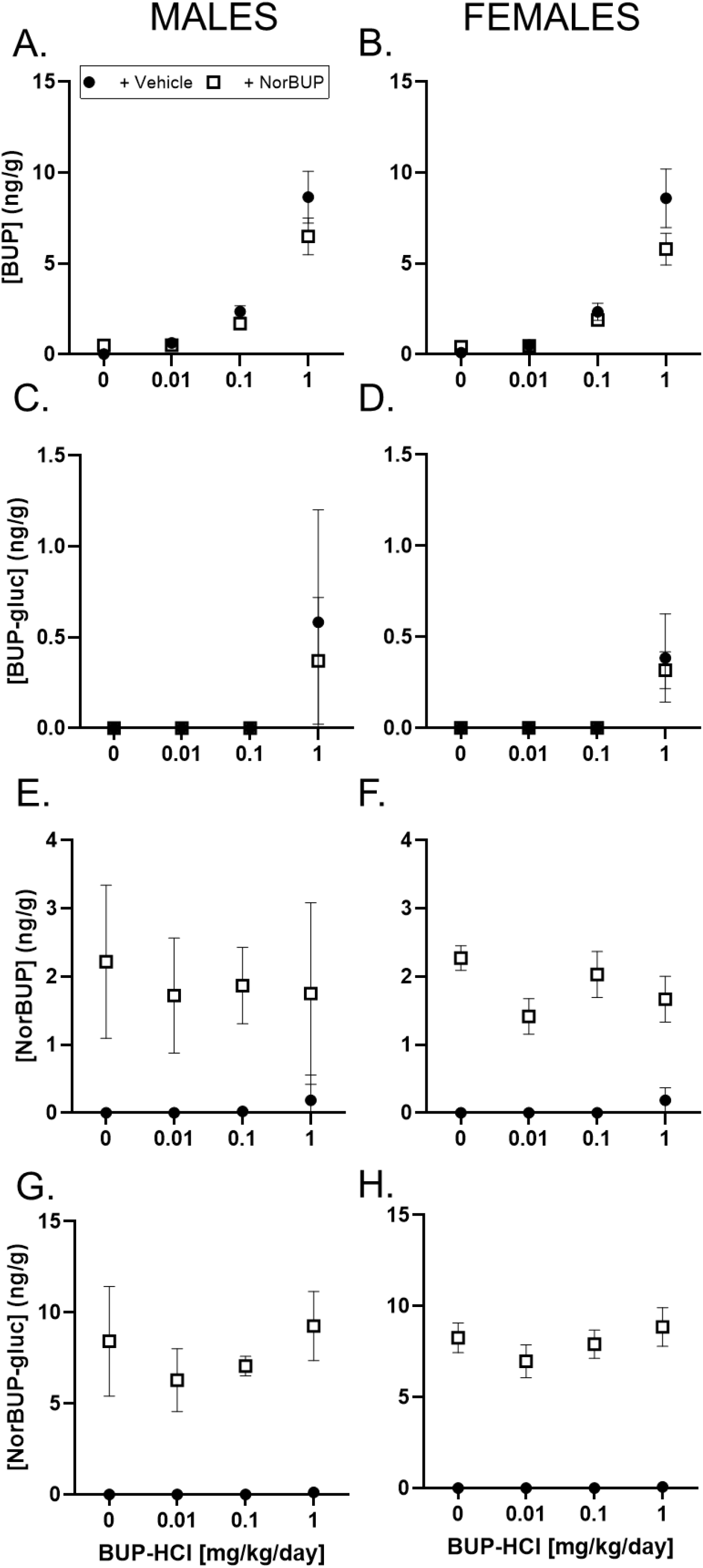
BUP, NorBUP, BUP-gluc and NorBUP-gluc metabolites were similar in both male and female neonatal brains. Neonatal male (A, C, E, and G) and female (B, D, F, and H) brain concentrations of BUP, NorBUP, BUP-gluc and NorBUP-gluc. Prenatal treatments are plotted on the x-axis and include NorBUP (1 mg/kg/day; s.c.), BUP (0.01, 0.1 and 1 mg/kg/day; s.c.) and vehicle (1:2:1 DMSO/PEG-400/saline, 0.120 mL/day, s.c.). Data points represented on the y-axis represent mean metabolite concentration. Error bars represent S.E.M. N = 4-7.

To investigate the relationship between NOWS and brain concentrations of BUP and NorBUP, we conducted multiple linear regression analyses modeling BUP and NorBUP concentrations in neonatal brains as predictors and movement duration following injection with the intermediate dose of NTX (1 mg/kg) as the dependent variable (or “outcome”). Separate analyses were conducted for males and females. For females, the model was significant [F (2, 40) = 23.97, *p<* 0.0001] and fit the data with an adjusted R^2^ = 0.52 (i.e., 52% of the variation in movement duration is explainable by the model). The equation for the model for females was Y = 27.14 + 10.62*BUP(β1) + 51.34*NorBUP(β2). The partial regression coefficients for BUP and NorBUP, but not the intercept, were significantly greater than zero (**Table 2**). For males, the model was also significant [F (2, 40) = 5.84, P = 0.0059] and fit the data with an adjusted R^2^ = 0.19 (i.e., 19% of the variation in movement duration is explainable by the model). The equation for the model for males was Y = 76.24 + 11.38*BUP(β1) + 19.21*NorBUP(β2). The intercept and partial regression coefficient for BUP, but not NorBUP, were significantly greater than zero (**Table 2**). For both males and females, the independent variables had low collinearity (VIF ≤1.012) and the models passed normality testing.

**Table 2.** Multiple linear regression model predicting movement duration (s)

|  |  |  |  |  |  |
| --- | --- | --- | --- | --- | --- |
| **Parameter estimates** | **Variable** | **Estimate** | **95% CI** | **\|t\|** | **p-value** |
| **FEMALE** |  |  |  |  |  |
| $\beta_0$ | Intercept | 27.14 | -3.73,<br>58.02 | 1.78 | .0832 |
| $\beta_1$ | [BUP]<sub>neonatal brain</sub> | 10.62 | 4.25,<br>16.98 | 3.37 | **.0017** |
| $\beta_2$ | [NorBUP]<sub>neonatal brain</sub> | 51.34 | 33.58,<br>69.09 | 5.84 | **<.0001** |
| **MALE** |  |  |  |  |  |
| $\beta_0$ | Intercept | 76.24 | 36.27,<br>116.2 | 3.85 | **.0004** |
| $\beta_1$ | [BUP]<sub>neonatal brain</sub> | 11.38 | 3.01,<br>19.75 | 2.75 | **.009** |
| $\beta_2$ | [NorBUP]<sub>neonatal brain</sub> | 19.21 | -3.320,<br>41.74 | 1.72 | .093 |

Lastly, we examined the interaction between BUP and NorBUP to activate hMORs by measuring [^35^S]GTPγS specific binding in CHO-hMOR homogenates in response to various concentrations of BUP (0, 3, 10, 30, 100, 300, and 1000 nM) in the absence or presence of NorBUP (0 or 1000 nM). As expected, BUP (3-1000 nM) and NorBUP (1000 nM) each independently increased [^35^S]GTPγS specific binding relative to vehicle-induced specific binding, with NorBUP exhibiting higher efficacy than BUP (**Fig 6**). BUP and NorBUP in combination significantly interacted (F [6, 24] = 327.7; *p<*0.0001) such that increasing BUP concentrations decreased NorBUP-induced receptor activation [IC^50^ (95%CI) = 58.73 nM (28.15, 132.0 nM)]. Only the highest BUP concentration, which was equimolar to NorBUP, fully antagonized NorBUP-induced specific binding to BUP-induced levels (“ns”, **Fig 6**). Lower BUP concentrations either partially antagonized (30-300 nM) or increased (3-10 nM) specific binding induced by NorBUP (1000 nM).

**Figure 6.**
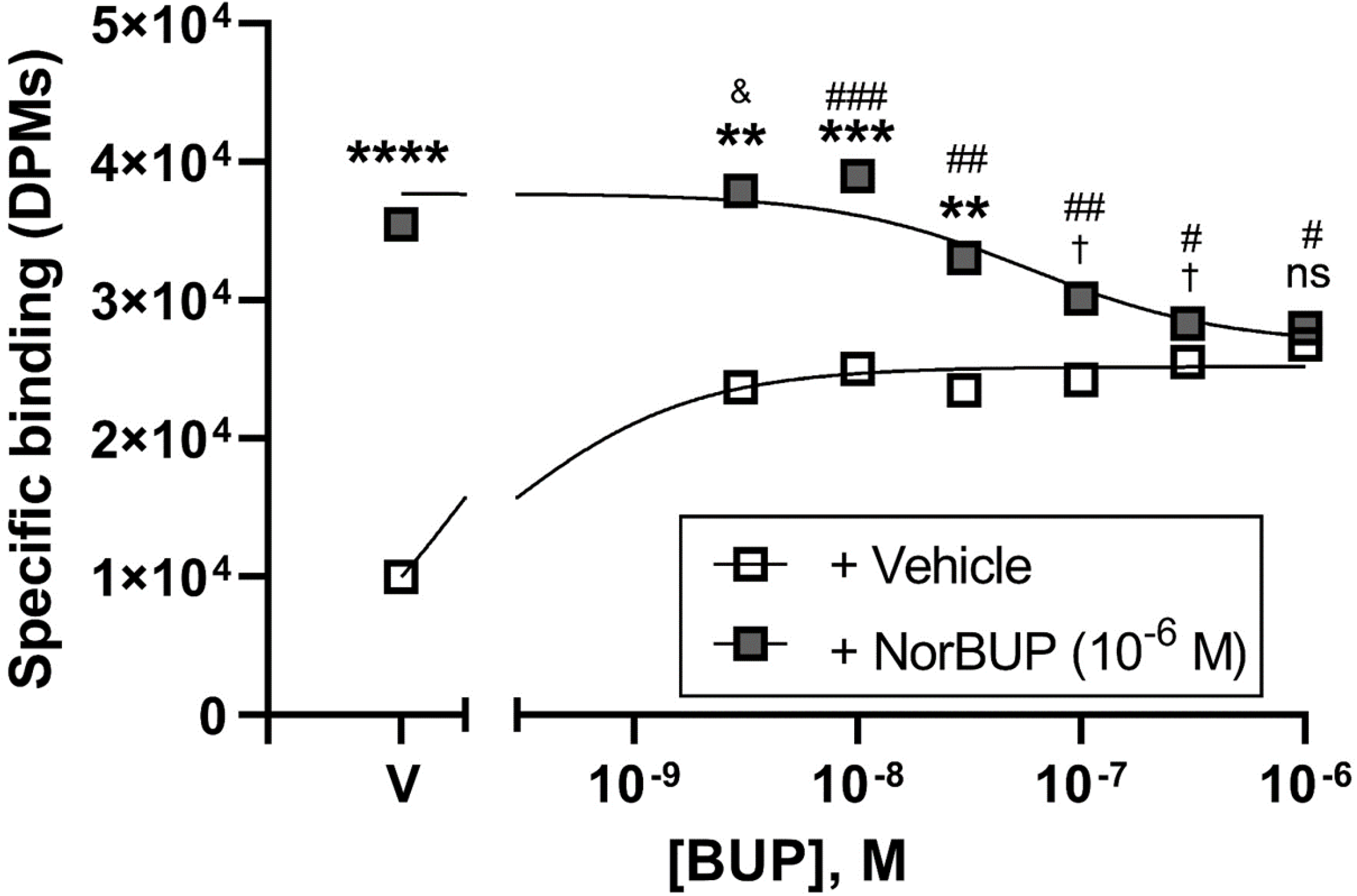
BUP dose-dependently antagonizes NorBUP-induced activation of hMOR. Activation of hMOR was measured as specific binding of [^35^S]GTPγS in decays per minute (DPMs, y-axis) as a function of BUP concentration (x-axis) in the presence or absence of 10 μM NorBUP (filled or open squares, respectively). Squares and error bars represent means and standard error of the mean of three independent experiments that were each performed in triplicate. ^†^P < 0.10, **P < 0.01, ***P < 0.001, ****P < 0.0001 vs BUP + vehicle, Sidek’s multiple comparisons test. ^&^P < 0.10, ^#^P < 0.05, ^##^P < 0.01, ^###^P < 0.001 vs NorBUP + vehicle.

## 4. Discussion

The mechanisms underlying BUP-associated NOWS are poorly understood. The present study is the first to model a wide range of clinically relevant BUP:NorBUP dose ratios (1:1, 10:1, and 1:100) and characterize the relationship between maternal dose, neonatal brain concentrations, and NOWS. Our two-part hypothesis was that BUP will not antagonize NorBUP-induced NOWS and that NorBUP co-administered with BUP will increase NOWS severity compared to BUP administration alone. We conclude that BUP did not antagonize NorBUP based on our observation that co-administering BUP and NorBUP did not reduce naltrexone-precipitated withdrawal severity relative to NorBUP alone. These findings are important because they suggest NorBUP can induce fetal opioid dependence even in the presence of BUP and thus provides more evidence implicating NorBUP in BUP-associated NOWS. Furthermore, 1 mg/kg/day BUP enhanced the effect of NorBUP on NOWS severity in females, but not males, that were challenged with 1 mg/kg NTX, suggesting that BUP and NorBUP are additive in a sex-dependent manner. We additionally determined that the relationship between NOWS and neonatal brain concentrations of NorBUP is sex-dependent such that NorBUP brain concentrations are more closely associated with NOWS in females than in males. Altogether, these findings importantly suggest that NorBUP can cause fetal opioid dependence, even in the presence of BUP, and that females are more sensitive than males to fetal opioid dependence caused by prenatal NorBUP exposure.

In our study, male and female neonates had similar brain concentrations of BUP, NorBUP, BUP-gluc, and NorBUP-gluc. This suggests that sex differences in the contribution of NorBUP to NOWS are likely mediated by sex differences in the pharmacodynamics of NorBUP in the fetal brain and not pharmacokinetics (e.g., sex differences in placental permeability or fetal metabolism). This implies that the fetal female brain is more sensitive than the fetal male brain to chronic NorBUP-induced changes in opioid receptor signaling at the cellular and sub-cellular levels. This interesting finding seems to contradict studies that found male babies experience more severe NOWS than female babies (O’Connor et al., 2013) and that male sex is an important risk factor for NOWS (Charles et al., 2017). However, these studies were not specific to BUP treatment, and BUP-specific studies have reported no sex effect of BUP on NOWS (Liu et al., 2010; Shah et al., 2016). The contribution of NorBUP to BUP-associated NOWS, possibly mediated by population variability in BUP metabolism to NorBUP (Meaden et al., 2021), may help explain the unpredictability of NOWS following prenatal BUP treatment (Bakstad et al., 2009; Jones et al., 2014; Kacinko et al., 2008), but NorBUP does not cause a clear sex difference in BUP-associated NOWS in humans. Ultimately, our finding that withdrawal is more heavily influenced by NorBUP in females than males suggests that strategies to reduce NorBUP formation and fetal NorBUP exposure may be more therapeutically beneficial for female fetuses than for male fetuses. Future studies that compare opioid receptor density and function (e.g., [^35^S]GTPγS binding in response to receptor-selective agonists) in brains from neonatal males and females (opioid-naïve and prenatally exposed to BUP and/or NorBUP) would help determine whether the sensitivity is mediated at the levels of the receptor. Interestingly, in both female and male neonatal brains, NorBUP-gluc concentrations were 3- to 4-fold higher than NorBUP concentrations, suggesting extensive maternal-fetal glucuronidation of NorBUP and sequestration of NorBUP-gluc in the neonatal brains. Fetal and neonatal glucuronidation capacity is very low (Allegaert et al., 2009), meaning that NorBUP-gluc in the fetal brain was most likely formed by the maternal system, crossed the placenta from maternal circulation, and accumulated in the fetal brain. Although NorBUP-gluc has low or no affinity for opioid receptors, a subcutaneous injection (2.2 mg/kg) markedly reduced the tidal volume of breathing and locomotor activity in mice (Brown et al., 2011), suggesting NorBUP-gluc has biological activity. The possibility that NorBUP-gluc has biological activity along with our finding of accumulation of NorBUP-gluc in the fetal/neonatal brain indicate that future studies are needed to investigate the effects of chronic fetal exposure to NorBUP-gluc.

In contrast to NorBUP-gluc, BUP-gluc concentrations in the neonatal brain were very low, despite BUP-gluc being structurally similar to NorBUP-gluc and presumably having similar physiochemical properties. Low neonatal brain concentrations of BUP-gluc is unlikely to be due to low maternal plasma concentrations of BUP-gluc because BUP is readily converted to BUP-gluc and plasma concentrations of the BUP-gluc and NorBUP-gluc are similar (Joshi et al., 2017)following administration of BUP to rats, suggesting factors other than maternal plasma concentration determine fetal and neonatal brain concentrations of BUP-gluc. However, one limitation of this study is the lack of maternal plasma sampling to quantify BUP, NorBUP, and their glucuronides to definitively relate maternal plasma concentrations to fetal brain exposure and NOWS. Another limitation is the analysis of neonatal brains instead of fetal brains, which would provide a more accurate snapshot of *in utero* exposure to the opioids and their metabolites. Because our primary endpoint of interest was neonatal opioid withdrawal, fetal brain harvest was precluded and we declined to sample maternal plasma to avoid potentially confounding the study by stressing the dams. Additional studies are needed to determine whether, under the experimental conditions used in this study, fetal brain concentrations replicate the neonatal brain concentrations we report in this study, and whether maternal plasma concentrations of BUP-gluc and NorBUP-gluc are similar. If BUP-gluc plasma concentrations are relatively similar to those of NorBUP-gluc, then there are likely different mechanisms or factors that control placental passage of BUP-gluc and NorBUP-gluc.

To better understand interactions between BUP and NorBUP at the receptor level, we measured hMOR-mediated G-protein activation by NorBUP in the presence of increasing concentrations of BUP. This approach allowed us to control concentrations of each drug at the receptor and directly measure receptor activation as a consequence of drug-receptor interactions in a manner that is not possible in whole-animal assays due to pharmacokinetic factors. With this assay, we determined that complete antagonism of NorBUP by BUP occurs only at BUP concentrations that are equal to NorBUP. This was unexpected because BUP has extremely high affinity for hMOR and slow dissociation from hMOR (Huang et al., 2001; Obeng et al., 2021); thus, we expected BUP to fully antagonize NorBUP at much lower BUP concentrations. These data are also interesting because low concentrations of BUP appeared to have additive effects on NorBUP-induced activation of hMOR that transitioned to antagonistic effects as BUP concentrations were increased. This suggests there are complex interactions between NorBUP and BUP at the level of the receptor. There are limitations to this experiment. There are still questions left to answer regarding the kinetics of the interaction between BUP and NorBUP because 1) only one incubation time point is represented and, 2) once a G-protein is activated and bound by [^35^S]GTPγS, the binding is irreversible it can no longer recycle to a deactivated state and re-activated. Future studies that vary the incubation time points, and/or withhold addition of the irreversible ligand [^35^S]GTPγS until after equilibrium of receptor binding by BUP and NorBUP is reached would be highly valuable for better understanding BUP and NorBUP interactions in the fetal brain.

Some findings relating to the neonatal withdrawal data were unexpected. In our rat model of NOWS, an intraperitoneal injection of saline is used as a control for stress and movement induced by handling and injection, as well as spontaneous withdrawals the pup may be undergoing. The saline injection, which generally elicits very little pup movement even in highly opioid-dependent pups (Griffin et al., 2019), enables assessment of antagonist-precipitated withdrawal severity by comparisons with behaviors elicited by injection of NTX. Intriguingly, prenatal exposure to 1 mg/kg/day BUP (independent of NorBUP exposure) robustly increased movement duration following injection with either saline or NTX. The magnitude of the increase was equal to that observed in pups prenatally exposed to NorBUP and challenged with a high dose of NTX, which is unexpected given the differences between BUP and NorBUP in MOR efficacy (i.e., BUP has low efficacy and NorBUP has high efficacy). Rat pups exhibiting spontaneous withdrawal signs only three hours after delivery would be surprising because neonatal onset of spontaneous withdrawal from buprenorphine in humans occurs approximately 40 hours post-delivery and peaks at 70 hours post-delivery, but can be delayed until 5 days or more (Hudak et al., 2012). Although the time course of spontaneous neonatal opioid withdrawal in rats has not been characterized, it is highly unlikely that the neonatal pups would display such robust spontaneous withdrawal behaviors within the first day of life. An alternative explanation of this phenomenon is opioid-induced hyperalgesia (Lee et al., 2011), a condition characterized by a paradoxical increase in perception of painful stimuli and nociceptive sensitization. For example, the intraperitoneal saline injection may be significantly more painful for pups with chronic prenatal exposure to a high dose of BUP. Chronic use of BUP has been reported to cause hyperalgesia resulting in exacerbation of pain sensation rather than its relief (Athanasos et al., 2019; Larson et al., 2020; Mercadante et al., 2019). Although opioid-induced hyperalgesia is generally associated with opioid use by adults, evidence of opioid-induced hyperalgesia in infants (Efune and Rebstock, 2022; Hallett and Chalkiadis, 2012; Zissen et al., 2007) has been reported and is a potentially under-recognized phenomenon due to difficulty distinguishing whether signs of pain and discomfort in non-verbal infants are due to opioid-induced hyperalgesia or inadequate pain management that is possibly caused by tolerance. Hyperalgesia is a symptom of opioid withdrawal, further complicating distinction between BUP-induced hyperalgesia and withdrawal in the neonatal human or rat. Future studies comparing nociceptive responses (e.g., movement duration) in BUP-dependent neonatal rats immediately after dosing versus those abstinent from buprenorphine for several hours (similar to pups in our experiment) would help distinguish BUP-induced hyperalgesia from withdrawal, since pups would not be in withdrawal immediately following BUP administration.

The most important findings of our study are that NorBUP can cause NOWS even in the presence of BUP and that NorBUP has more influence in BUP-associated NOWS in females than in males. This provides a strong rationale for strategies that reduce fetal NorBUP exposure by reducing BUP metabolism to NorBUP (Janganati et al., 2022), and thus are likely to decrease the incidence and severity of NOWS.

## Abbreviations

ANOVA: analysis of variance
BUP: buprenorphine
BUP-gluc: buprenorphine glucuronide
DMSO: dimethyl sulfoxide
GD: gestational day
LC-MS-MS: liquid chromatography-tandem mass spectrometry
NTX: naltrexone
NAS: neonatal abstinence syndrome
NorBUP: norbuprenorphine
NorBUP-gluc: norbuprenorphine glucuronide
NOWS: neonatal opioid withdrawal syndrome
OUD: opioid use disorder
PEG-400: polyethylene glycol-400.

## Funding sources

This project is supported by the Translational Research Institute (TRI), grants UL1 TR003107 and TL1 TR003109 through the National Center for Advancing Translational Sciences of the National Institutes of Health (NIH) and by a UAMS Barton Pilot Study Award. Buprenorphine HCl and norbuprenorphine were provided by the NIDA Drug Supply Program. The funding sources had no involvement in the preparation of the manuscript or the decision to submit it for peer review and publication.

## Role of Funding Source

The funding sources had no involvement in the preparation of the manuscript or the decision to submit it for peer review and publication.

## Contributors

Conception and design: JT, MDB, LKB; Acquisition of Data: JT, PS, BJP, LUC, LKB; Analysis and interpretation of data: JT, PS, BJP, MDB, LUC, GSG, LKB; Drafting the manuscript: JT, LKB; Revising the manuscript: All authors; Approval of the final article: All authors.

## Conflict of Interest

No conflict declared

## Acknowledgements

This project is supported by the Translational Research Institute (TRI), grants UL1 TR003107 and TL1 TR003109 through the National Center for Advancing Translational Sciences of the National Institutes of Health (NIH) and by a UAMS Barton Pilot Study Award. Buprenorphine HCl and norbuprenorphine were provided by the NIDA Drug Supply Program.

## Notes

### Competing Interest Statement

The authors have declared no competing interest.

